# Evolution of the olfactory system during the radiation of Heliconiini butterflies

**DOI:** 10.1101/2025.04.06.647427

**Authors:** Yi Peng Toh, Francesco Cicconardi, Giorgio Bianchini, Richard M Merrill, Stephen H. Montgomery

**Affiliations:** Division of Evolutionary Biology, Faculty of Biology, LMU Munich, Germany; School of Biological Science, University of Bristol, Bristol, United Kingdom; School of Geographical Sciences, University of Bristol, Bristol, United Kingdom

**Keywords:** olfactory system evolution, comparative neuroanatomy, Heliconiini, developmental plasticity, antennal lobe morphology

## Abstract

Sensory system evolution plays a crucial role in shaping species’ interactions with their environment, yet the extent to which olfactory system diversity reflects ecological and evolutionary pressures at a macroevolutionary scale remains unclear. Here, we investigate the evolution of the olfactory system across the Heliconiini butterfly tribe, an ecologically diverse but closely related group. Using a comparative approach, we examined variation in antennal lobe morphology and its constituent structures, the glomeruli and antennal lobe hub, as well as olfactory receptor repertoires across species. We found that antennal lobe size variation is driven by independent shifts in glomerular and antennal lobe hub volumes, with species-specific differences occurring against a backdrop of broader phylogenetic stability. While no direct associations with ecological traits were observed, certain species showed large expansions in total glomerular volume and olfactory receptor numbers, warranting further investigation into unmeasured ecological or behavioural factors. Additionally, comparisons between wild-caught and insectary-reared individuals revealed a surprising pattern of developmental plasticity, with antennal lobe hub volumes increasing and glomeruli volumes decreasing in captivity, highlighting the influence of environmental conditions on neural development. These findings suggest that olfactory evolution in Heliconiini is shaped by both evolutionary divergence and developmental plasticity, emphasizing the need to integrate phylogenetic, ecological, and developmental perspectives to fully understand sensory system adaptation.

## Introduction

When confronted with new or changing environments, populations must adapt to different selective pressures (Darwin, 1859; Coyne & Orr, 2004; Rundle & Nosil, 2005; Schluter, 2009). These adaptations are often behavioural, which ultimately depend on changes in sensory and/or neural systems (Endler, 1992; Boughman, 2002; Montgomery et al., 2021). As such, sensory system evolution allows organisms to meet challenges imposed by their environment by increasing their ability to detect and respond to environmental cues (Dell’Aglio et al., 2024a). For example, cichlids living in depth-differentiated environments display shifts in sensitivity towards long and short-wavelength light, which are associated with habitat-dependent survival (Wright et al.,2019, Seehausen et al., 2008; Maan et al., 2006), and in New World warblers, differences in opsin gene expression are similarly associated with habitat (Bloch, 2015). Beyond the visual system, evidence also exists for adaptive divergence of the mechanoreception, electroreception and olfactory systems (Morris et al., 2021; Arnegard et al., 2010; Wark & Peichel, 2010).

The olfactory system plays a crucial role in most animals, as odour detection is often critical for both mate-finding and foraging (Togunov et al., 2017; Zjacic & Scholz, 2022). In insects, the primary olfactory processing centre is comprised of paired sphere-shaped structures within the midbrain called the antennal lobes, analogous to the olfactory bulbs in mammals (Boeckh et al., 1990) (Fig. 1). Each antennal lobe is made up of glomeruli positioned around a central axonal network of mainly olfactory projection neurons and local interneurons, called the antennal lobe hub (Malnic et al., 1999; Montgomery et al., 2016; Kymre et al., 2021). Each glomerulus houses the terminal projections of olfactory receptor neurons and the dendritic arbors of local interneurons, which help to arbitrate complex interactions across the antennal lobe (Fusca & Kloppenburg, 2021). Different odours are recognized by different combinations of olfactory receptors, and this translates to different glomeruli being activated, such that olfactory information is represented by a combinatorial coding strategy (Hansson et al., 1992; Malnic et al., 1999; Hallem & Carlson, 2006; Bisch-Knaden et al., 2018). This activation pattern is then relayed via projection neurons, which also synapse in the glomeruli, to integrative processing centres in the brain, the mushroom bodies and the lateral horn (Homberg et al., 1989) (Fig 1).

**Figure 1:**
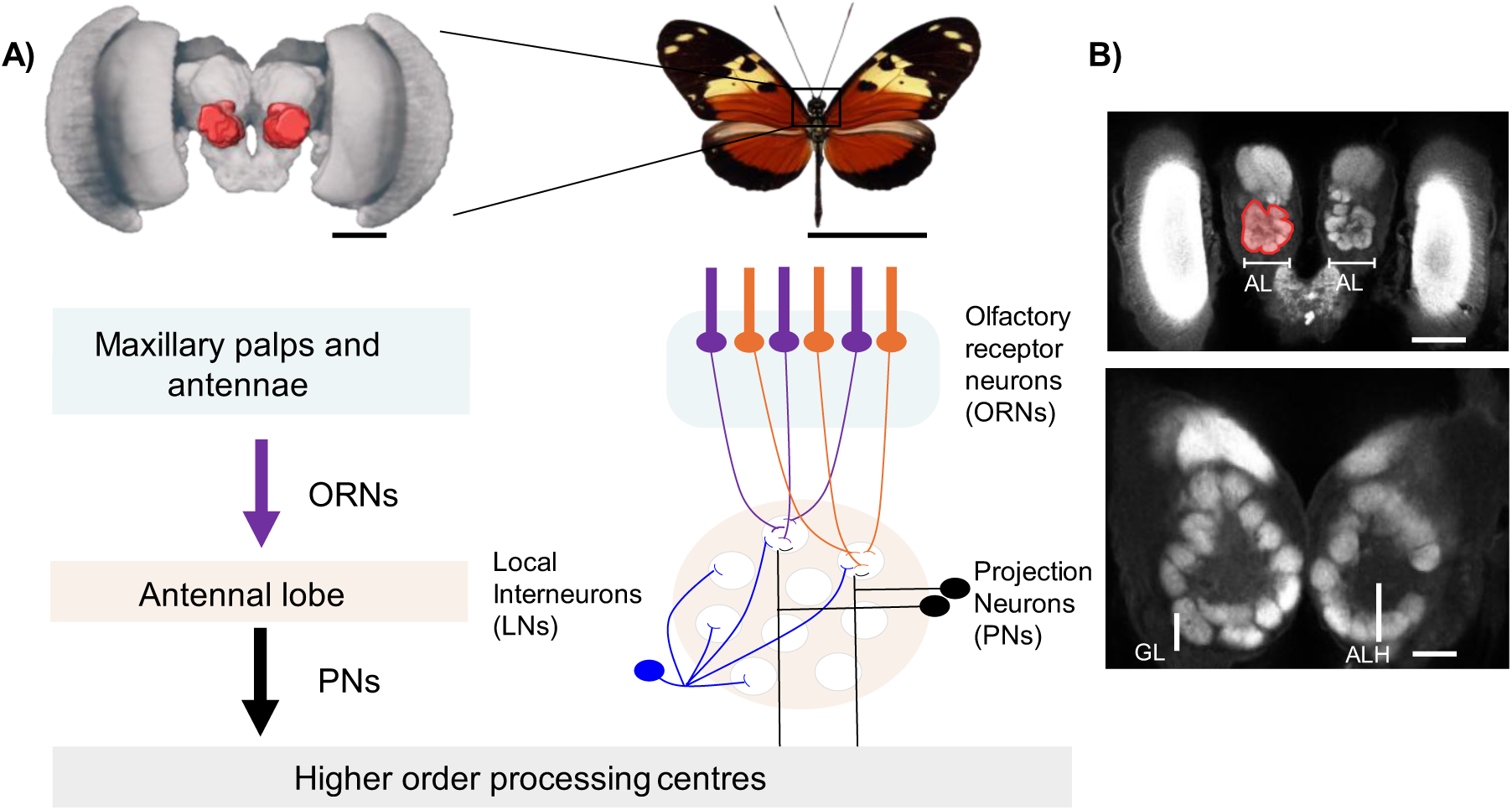
Olfactory pathway in invertebrates, including an overview of the antennal lobes and their constituent components in the brain. **A)** Odours are detected by the olfactory receptor neurons (ORNs) located in the maxillary palps and antennae in invertebrates. Sensory information is then passed on by the ORNs to the first processing centre in the brain, the antennal lobe (AL). This centre is made up of glomeruli (GL) and antennal lobe hub (ALH) which houses several different types of neurons, ORNs, local interneurons (LNs) and projection neurons (PNs). LNs help to arbitrate complex interactions in the antennal lobe. The signal is then forwarded to higher order processing centres by the PNs. The butterfly used in the figure is *H. hecale*. Figure concept adapted from Norwegian University of Science and Technology; https://shorturl.at/JAiDS. Scale bar: rendered volume of whole brain: 500μm; *H. hecale*: 25mm **B)** Top: view of the antennal lobes in the X-Y plane of a 3D image confocal stack of a Heliconiini brain; Bottom: zoomed-in view of the glomeruli and antennal lobe hub, the constituent units of the antennal lobe. Scale bar: Top: 500μm, Bottom: 50μm.

Individual olfactory receptor neurons originate from peripheral structures, predominantly the antennae, but also the maxillary palps and legs, and terminate within one of the antennal lobe glomeruli (Briscoe et al., 2013; Hansson & Anton, 2000; Arnold et al., 1985; Boeckh et al., 1977; Homberg et al., 1989). Two general principles of olfactory system architecture are that each olfactory receptor neuron only expresses a single olfactory receptor, and that each glomerulus receives input from only one class of olfactory receptor neurons. This essentially results in a one-receptor-one-glomerulus relationship (Hildebrand & Shepherd, 1997). Although this paradigm has been recently challenged by evidence of co-expression of multiple receptors in single receptor neurons in *Aedes aegypti* and *Drosophila melanogaster* (Herre et al., 2022; Task et al., 2022), it remains unclear how taxonomically widespread this exception is.

Insect olfactory receptors function as heteromeric tetramers, consisting of a single ligand-binding olfactory receptor and a coreceptor, *Orco* (Störtkuhl and Kettler, 2001; Dobritsa et al., 2003; Vosshall et al., 2000; Larsson et al., 2004). Except for *Orco*, the conservation of close orthologs and subfamily structures are not detected across insect orders (Hansson & Stensmyr, 2011). Studies looking at the number of olfactory receptors in invertebrates range widely from around 10 in Phthiraptera (i.e., lice (Kirkness et al., 2010)) to ∼200–400 in Hymenoptera (i.e., bees, ants, and wasps (Robertson et al., 2010, Wurm et al., 2011)), suggesting olfactory receptor repertoires are both highly variable across species and intimately linked to species’ ecology. Among closely related species, olfactory receptors show a high rate of birth and death, even when overall receptor numbers remain relatively constant (Cicconardi et al., 2024; Nei & Rooney, 2005) and are often associated with signatures of positive selection (Rooney et al., 2002; Piontkivska et al., 2002; Cicconardi et al., 2017). This rapid turnover suggests that olfactory receptor repertoires can evolve quickly to meet immediate ecological demands. For example, comparisons amongst close relatives in *Drosophila* showed an increased expression and upregulation of odorant receptor and odorant-binding proteins in ecological specialists compared to more generalist species (Kopp et al., 2008). As another example, in closely related species of *Heliconius* butterflies living in divergent environments, widespread gene expression divergence was also found, consistent with evolutionary divergence in the sensory tissues, and this was argued to have facilitated speciation in the early stages (Wu et al., 2022). Thus, together with the broader loss of conserved subfamilies across insect orders indicating deep evolutionary divergence over millions of years, these examples suggest dynamic patterns of evolution in olfactory receptor repertoires across both long and short evolutionary timescales (Yohe et al., 2020).

While most research on sensory system evolution has focused on changes in these olfactory receptor gene repertoires (e.g. Carlsson et al., 2013; Maan et al., 2006; Bloch, 2015; Wark & Peichel, 2010), variation in downstream neural processing also plays a significant role in olfactory evolution. Comparative studies often reveal volumetric differences in regions of the brain involved in sensory processing, which likely reflects changes in cell number or size (Gronenberg & Hölldobler, 1999; Kondoh et al., 2003; Streinzer et al., 2013; O’Donnell et al., 2013; Montgomery & Ott, 2015). For example, detailed studies in *Drosophila* have not only revealed evolution in the number of olfactory receptor genes and the types of olfactory sensilla, but also volumetric changes in the antennal lobe and specific glomeruli (Auer et al., 2020; Prieto-Godino et al., 2020, 2021; Linz et al., 2013). This provides evidence that olfactory evolution can occur at multiple points along the olfactory pathway and illustrate the advantages of studying olfactory system evolution in the context of an adaptive radiation, where closely related species are ecologically distinct (Depetris-Chauvin et al., 2023; Auer et al., 2020; Prieto-Godino et al., 2016, 2017).

Studies of Lepidoptera are proving informative as case studies linking ecological divergence, behaviour and sensory evolution (Carlsson et al., 2013; Montgomery & Merrill, 2017; Montgomery et al., 2021; Hebberecht et al., 2023, Borrero et al., 2024). For example, differences in the overall volume of the antennal lobe have been found between nocturnal and diurnal moth species, with these volumetric differences corresponding to differences in sensory weighting during foraging experiments (Stöckl et al., 2016), lending support to the prevailing theory that expansions of certain brain regions correspond to enhanced information processing (Barton, 1998; Barton et al., 1995). Classic comparative studies in moths have also revealed differences in the arrangement and volume of a region of the glomeruli termed the macroglomerular complex (Boeckh and Boeckh, 1979; Hansson et al., 1991; Namiki et al., 2014; Berg et al., 2014; Liu et al., 2021). The macroglomerular complex is enlarged in males (or is male specific) and is involved in receiving and processing information about female sex pheromones (Koontz and Schneider, 1987). In contrast to moths, however, most butterflies lack a macroglomerular complex (Rospars, 1983; Heinze and Reppert, 2012; Carlsson et al., 2013), although a similar cluster of morphologically distinct glomeruli has re-emerged at least once in butterflies, in the tribe Ithomini, associated with increased expression of a small number of olfactory receptors (Montgomery & Ott, 2015; Morris et al., 2021; Cicconardi et al., 2024).

The Nymphalid butterfly tribe, Heliconiini, includes the diverse genus *Heliconius*, which has radiated across the Neotropics to now include ∼50 species, having diverged from its sister genus *Eueides* as recently as 18 million years ago (Kozak et al., 2015, Cicconardi et al., 2023). Although *Heliconius* research has largely focused on visual cues and behaviour (e.g. Dell’Aglio et al., 2016; Rossi et al., 2024, Merrill et al., 2015, Hausmann et al., 2023), chemoreception plays an important role in many aspects of *Heliconius* behaviour. *Heliconius* males produces sex and species-specific compounds which alter female mating decisions (Darragh et al., 2017; Mérot et al., 2015, González-Rojas et al.,2020), and are also known to generate anti-aphrodisiac pheromones, which are transferred to females during copulation, and repel subsequent males (Schulz et al., 2008; Gilbert, 1976; Estrada et al., 2011). Differences in olfactory receptor expression have also been documented between the sympatric species, *H. cydno* and *H. melpomene*, which have distinct mating and host plant preferences, highlighting the potential for olfactory evolution in maintaining species barriers in this group (Darragh et al., 2017; van Schooten et al., 2020). The antennal lobe also appears to show frequent differences in size, possibly associated with habitat shifts, between closely related *Heliconius* species (Montgomery & Merrill, 2017; Montgomery et al., 2021; Hebberecht et al., 2023). However, while these studies have provided valuable insights, they have largely been confined to comparisons between individual taxon pairs, leaving us without a comprehensive understanding of chemosensory evolution across the Heliconiini.

To further investigate what drives changes in the olfactory system across the Heliconiini, we expanded neuroanatomical and molecular comparisons across the whole Heliconiini tribe. Specifically, we explored how the volume of the antennal lobe, as well as its constituent units, the antennal lobe hub and glomeruli, in addition to the number of olfactory receptors, vary across the Heliconiini. We tested whether these components of the olfactory system are associated with ecological variation. Finally, we made use of data from individuals reared under common garden conditions to determine if any of the morphological variation we observed across wild populations can be attributed to developmentally induced plasticity.

## Methods

### Sampling across Heliconiini

We took advantage of an extensive collection of brain samples previously used to examine mushroom body evolution across the Heliconiini (Couto et al., 2023). These include 318 wild butterflies from 41 species (average 8 individuals per species; Fig. 2) belonging to the Heliconiini tribe sampled across sites in Costa Rica, Panama, French Guiana, Ecuador and Peru. We also made use of total antennal lobe volume segmented in prior studies (Couto et al., 2023; Hebberecht et al., 2023) but extended these data by re-segmenting these samples to include total antennal lobe, total glomeruli and antennal lobe hub volume.

**Figure 2:**
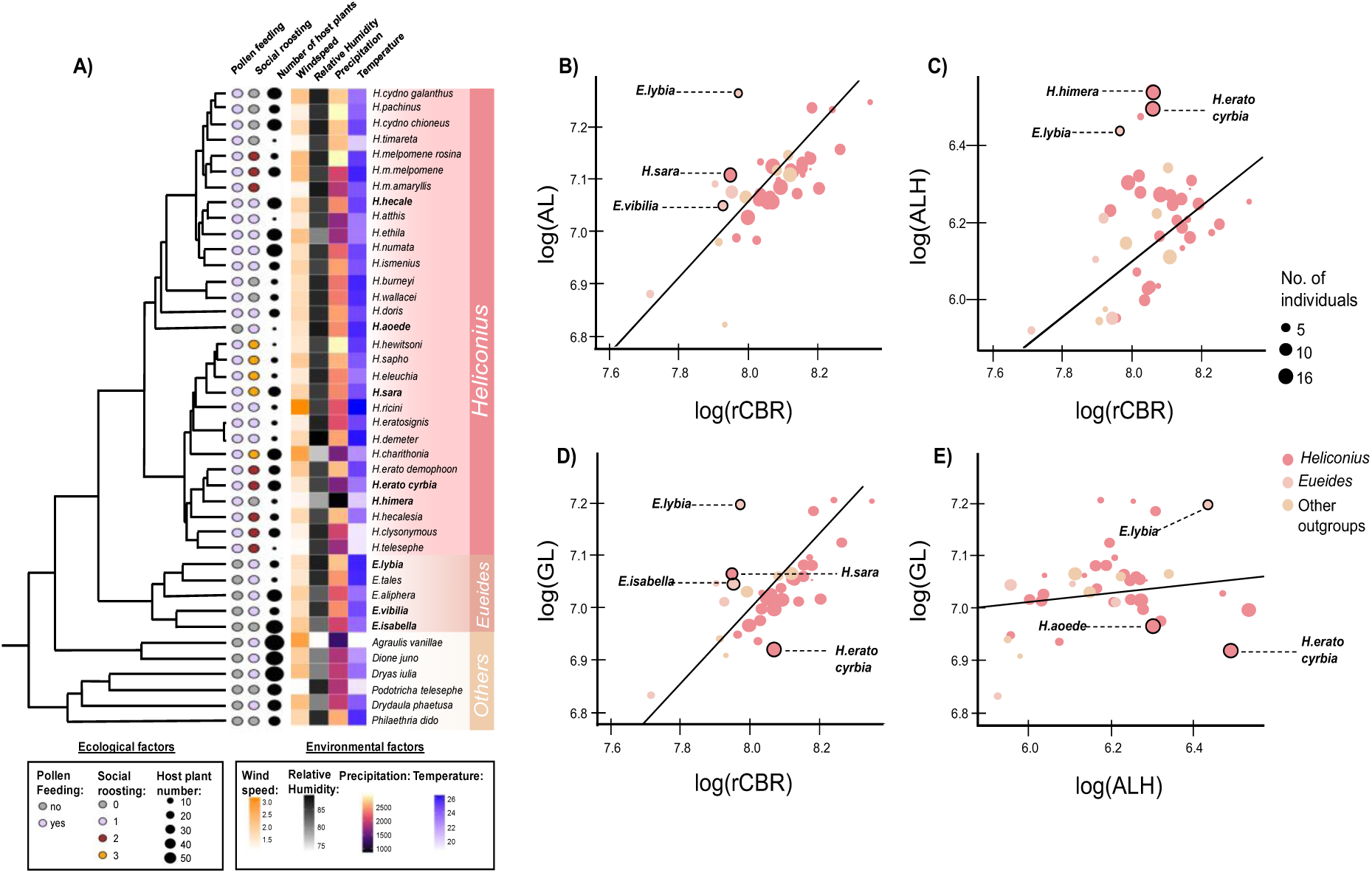
Variation in the size of olfactory neuropils and associated ecological and environmental traits in the Heliconiini. **A)** Phylogeny showing the Heliconiini species sampled and the major phylogenetic groupings that were also used in the comparative analyses. Species that displayed the greatest number of pairwise differences in terms of AL, ALH and GL volume, as well as the change in GL∼ALH volume are in bold. The associated environmental and ecological traits for each species are plotted at the tips of the tree. **B)** Variation in the AL volume in the Heliconiini. **C)** Variation in the ALH volume in the Heliconiini. **D)** Variation in the GL volume in the Heliconiini. **E)** Variation in the GL∼ALH volume in the Heliconiini. The volumes were allometrically controlled by plotting them against the volume of the rCBR. The top 3 species that had the greatest number of pairwise differences in each comparison are circled and labelled in the plots (note there are 4 species labelled in D) as 2 species had a similar number of pairwise differences. Point size based on the number of individuals for each species.

To determine whether any variation we observed was due to environmentally induced plasticity, we additionally examined neuropil volumes for five *Heliconius* species raised under common garden conditions, as described in Hebberecht *et al*. (2023). In brief, stocks of *H. erato*, *H. melpomene*, *H. cydno*, *H. hecale* and *H. ismenius* were established from wild individuals caught in Gamboa and the nearby Soberanía National Park, Panama, and maintained in insectaries (c. 2 × 2 × 2 m) at the Smithsonian Tropical Research Institute. All butterflies were provided with a ∼10% sugar solution, and either *Lantana* spp. or *Psiguria* spp. flowers as a source of pollen. *Passiflora* spp. were provided for oviposition, and caterpillars were raised on their native host plant (Merrill et al., 2013). Adults were maintained for 2-3 weeks to ensure maturity before sampling, and their age recorded.

All brains had been dissected, imaged, and partially segmented for previous studies (Couto et al. 2023; Hebberecht et al., 2023). This followed established protocols that stain against a synaptic protein, illuminating synapse-dense regions of the brain referred to as neuropils (Ott, 2008). We additionally scanned one side of the antennal lobe, chosen at random, of two individuals of *E. lybia,* and three individuals of *E. aliphera* for imaging at higher resolution with an x-y resolution of 1024 × 1024 pixels using a Leica SP8, 10X 0.4NA lens, and the same settings as previously collected data. These higher-resolution antennal lobe stacks, together with original confocal scans of *Dryadula phaetusa* in the resolution of 512 x 512 pixels, were used to estimate the number of glomeruli present in these species by segmenting all observable glomeruli to complement prior estimations of glomeruli numbers for *H. hecale* and *H. erato* (Montgomery et al., 2016). Measurement error due to compression of neighbouring glomeruli blurring boundaries between individual glomeruli creates uncertainty in the precise count of the total glomeruli for an individual, leading to possible slight over, or under, estimation.

### Image segmentations

We reconstructed the antennal lobe into the antennal lobe hub and glomeruli in a total of 318 wild-caught individuals, representing 41 species in the Heliconiini tribe, as well as an additional 44 *Heliconius* insectary reared individuals. To do so, we manually segmented each neuropil in the whole brain based on the boundaries that we see using the *labelfield* module of AMIRA using the *magic wand* function. We then obtained the volume of each neuropil using the *material statistics module* in AMIRA. We reconstructed these neuropils on one side of the brain, chosen at random, and multiplied them by two to get the total volume across the brain. We used the volume of the rest of the central brain (rCBR) as an allometric control which was obtained by previous reconstructions of the constituent brain regions (Couto et al., 2023; Hebberecht et al., 2022).

### Ecological data

We obtained data for three ecological traits for each Heliconiini species, in addition to a number of climatic traits. The ecological traits were the presence/absence of pollen feeding, the number of host plant species exploited, and the degree of social roosting. Pollen feeding was categorised as a binary trait (present/absent) following Couto et al. (2023), host plant use number was obtained from McLellan and Montgomery (2024), and roosting behaviour was classified as either solitary, loose group, small group, or large group, which was factored respectively into levels 1-4, following Brown (1981). These ecological traits were chosen as possible factors that would affect the evolution of sensory systems in Heliconiini: first, pollen feeding, the active collection of pollen grains as an adult source of amino acids, may involve processing of floral cues (Gilbert, 1972; Young & Montgomery, 2020); second, host plant generalism potentially requires recognition of more olfactory cues compared to species that utilise only a small number of host plant species (Wang et al., 2022; Burger et al., 2013), as recognition of each specific host plant involves the detection of its unique blend of volatile organic compounds (Bruce & Pickett, 2011); third, some *Heliconius* species are known to roost together in large groups at night, potentially representing a site of high-density social behaviours (Mallet, 1986).

To obtain climatic data, we followed a previous protocol by Rivas-Sánchez et al. (2023). In brief, we first obtained geo-referenced occurrence data and associated climatic data for all 41 species from Rosser et al. (2012). Outliers above the elevational range of 2100m were excluded from the dataset. We took into account sampling biases by spatially thinning occurrence records with the thin function in *spThin* (Aiello-Lammens et al., 2015; distance parameter = 5 km). For each occurrence data point, we then extracted the climatic variables BIO1 (Annual Mean Temperature) and BIO12 (Annual Precipitation) (30-arc sec resolution [∼1 km2]) from WorldClim (Fick & Hijmans, 2017) as well as the variables, relative humidity and average surface wind speed (30-arc sec resolution [∼1 km2]) in CHELSEA (Karger et al., 2017). We took the mean values of each variable for each species in the thinned data set and used them as the environmental data for our individuals, meaning that individuals from the same species have the same value for the same environmental factor. We selected these traits as temperature, precipitation, relative humidity and wind speed have all been shown to affect the concentration of odourants in the air (Hill & Petrucci, 1999; Martin et al., 2011; Dalton, 2000; Kuehn et al., 2008; Nagappan et al., 2017). At higher temperatures, vapor pressure is increased thereby increasing the rate of diffusion of odourant molecules (Hill & Petrucci, 1999). Both temperature and high odour concentrations can impact neuronal activity (Martin et al., 2011; Dalton, 2000; Devaud et al., 2001; Devaud, 2003; Störtkuhl et al., 1999). Increased humidity may increase the capacity of the air to carry odourants and possibly make detection easier (Kuehn et al., 2008). An increase in precipitation would also increase humidity and so we included it in our analysis. Finally, wind speeds have also been reported as a potential environmental factor impacting odour reception and locating (Nagappan et al., 2017).

### Odorant receptor annotation, phylogeny, and orthology assignment

To obtain the olfactory receptor repertoire across Heliconiini, annotated olfactory receptor coordinates and protein sequences obtained from previous studies (Cicconardi et al., 2023, 2024) were used to extend the list of annotated genes for Heliconinae. To do so, we implemented a modification of the pipeline used in Cicconardi et al. (2023), which used a combination of manual and automatic procedures. In brief, we utilized Miniprot (Li, 2023) to map protein sequences onto the genomes of 72 species: 44 *Heliconius* species, 6 *Eueides* species, 8 species sister to *Heliconius* and *Eueides* within the Heliconiini tribe, 2 *Argynnini* species, and 12 outgroup species from the Nymphalid family (including those most distantly related to the Heliconinae). This comprehensive mapping aimed to infer orthologous relationships across the Nymphalid family. Coding sequences (CDS) were extracted from all mapping hits, and a number of features were computed: the conserved PFAM v33.1 domains, using HMMscan (Eddy 2011) and DAMA (Bernardes et al. 2015), the presence of the peptide signal, using SignalP v5.0b (Emanuelsson et al. 2007), and the R implementation of TMHMM (Krogh et al. 2001) to predict the number of transmembrane helices (TMHs). At each locus, the best annotation was therefore selected based on the optimal protein length, conserved domain length and score, presence of P-signal and best number of TMHs. The amino acid sequences were then aligned using CLUSTALW v2.1 (settings: dnamatrix = IUB; gapopen = 10; gapext = 0.1; gapdist = 10; iteration = TREE; numiter = 1000; clustering = NJ), and the phylogeny was inferred using approximate Maximum Likelihood search as implemented in FastTree v2.1.11 SSE3, using Le-Gascuel (2008) model with pseudocounts and the slow exhaustive search algorithm to search for neighbor-joining (Price et al. 2010). To root the tree, ORco was used as an outgroup, and gene orthology was subsequently assigned based on the phylogenetic tree or reference genes in Cicconardi et al. (2024). For the graphical representation of the phylogenetic tree, we adopted TreeViewer (Bianchini & Sánchez-Baracaldo 2024). Due to the large number of sequences, we collapsed all monophyletic groups of orthologous genes, showing their relationships, while the full phylogenetic tree is available in the supplementary materials (Figure S1).

### Statistics

All neuropil volumes were log_10_-transformed prior to analysis, and all analyses were run in R (v4.4.1; R Core Team, 2024). Using a recent phylogenetic tree of the Heliconiini (Couto et al., 2023) to control for phylogenetic effects, we ran a series of phylogenetic generalized linear mixed models (GLMM) with Gaussian distributions using the R package MCMCglmm v2.32 (Hadfield, 2010). We assigned species as a random effect and tested whether the scaling relationship between the i) the total antennal lobe volume, ii) antennal lobe hub volume, and iii) the total volume of the glomeruli, and the volume of the rest of the central brain (rCBR) changes across broad phylogenetic groups within the Heliconiini. Specifically, we tested for differences between a) *Heliconius* and non-*Heliconius* Heliconiini, b) between Heliconiini genera, and c) between sub-clades within *Heliconius* (excluding outgroups), including the effects of sex. We also ran a fourth model to test for shifts in the scaling relationship between the glomeruli and the antennal lobe hub across the three different phylogenetic groupings noted above. To determine to what extent the volumetric differences observed can be accounted for by phylogeny, we additionally calculated the posterior mean of Pagel’s lambda (λ), after running a null GLMM model, with no other variables other than the allometric control rCBR or antennal lobe hub. We subsequently conducted Likelihood ratio testing of a model with the observed lambda and a model that assumes phylogenetic independence, as described in Münkemüller et al. (2012).

To test for species-specific differences, we also conducted a pairwise comparison of the 29 species with more than 5 individuals using the posterior distribution of the random effects. Here we adjusted the significant threshold to 0.99 to account for false positives when conducting multiple pairwise comparisons. Finally, we explored associations between the antennal lobe, antennal lobe hub, and glomeruli (after correcting for allometry), as well as the scaling relationship between antennal lobe hub and glomeruli volume, with environmental and life history variables. Associations between the neuropil with environmental and ecological factors were tested separately by taking the best fitting of these MCMCglmm models and then including these factors individually. Throughout, model checking for convergence was done using the *gelman.diag* function and for auto-correlation using the *autocorr* function in MCMCglmm, in addition to visually inspecting the trace plots. All models were run for 500,000 iterations, with a burn-in of 10,000 and a thinning factor of 500.

We repeated the above analyses to i) test for an association between the number of olfactory receptors annotated for each Heliconiini species and the same three neuropils (antennal lobe, antennal lobe hub and glomeruli), and ii) test for association between the number of olfactory receptors and environmental and ecological factors. We referenced the olfactory receptor data from a previous study (Cicconardi et al. 2023) and controlled for the quality of the annotation by including the N50 value. We also repeated the analyses with an alternative control, specifically the Benchmarking Universal Single-Copy Orthologues (BUSCO) score (Simão et al., 2015). When using the BUSCO as the alternative control, we included only species with a BUSCO completeness score of more than 90%. All N50 values, BUSCO scores and BUSCO completeness score were similarly referenced from Cicconardi et al. (2023).

To test for potential developmental plasticity, we compared the volume of the total antennal lobe, the antennal lobe hub, and glomeruli, between wild-caught and insectary-reared individuals for 5 *Heliconius* species (*H. erato cyrbia, H. melpomene, H. ismenius, H. cydno* and *H. hecale*). We first tested for differences using a series of nested linear mixed models implemented in the R package lme4 (Bates et al., 2015). In each case, log_10_ transformed values were used as the dependent variable. We included the rest of the central brain (rCBR), species and origin (wild-caught vs insectary-reared) as fixed factors. We also included an interaction term between species and origin in our full model. Sex was included as a random factor in all models. To acquire the minimum adequate model (MAM), and to determine the significance of individual terms, we used Likelihood Ratio Tests. All p-values were subsequently adjusted with Bonferroni correction to account for multiple testing.

We next conducted standardized major axis regression analyses using the SMATR package in R (Warton et al., 2012) to identify specific pairwise species differences in the volume of the antennal lobe, the antennal lobe hub and glomeruli, as well as differences the scaling relationship between antennal lobe hub and glomeruli volume in wild-caught and insectary-reared individuals. Following the allometric relationship *y*=*β*log *x* + *α,* we first tested for significant shift in slope (*β*), which would suggest a change in the allometric scaling exponent between the groups (Kruska, 2005). If a common slope was found, we subsequently tested for a shift in elevation (*α*), or a major-axis shift. Elevation shifts (or “grade-shifts”) suggest non-allometric divergence with a change in the intercept (Kruska, 2005), while a major-axis shift suggests that the groups change along a shared allometry which underlies changes in neuropil volume in concert with the rest of the brain (Barton & Harvey, 2000; Montgomery et al., 2016). Correction for multiple comparisons was done using a Sidak-correction, available as an option in the *sma* function of the SMATR package.

## 3. Results

### Variation in antennal lobe investment is driven by species-level differences rather than broad phylogenetic trends

Across all individuals, we saw considerable variation in the raw volumes of the antennal lobe (AL), which varied 6.2-fold across Heliconiini (from ∼3.8×10^6^μm^3^ to ∼23.5×10^6^μm^3^). This was reflected in variation in both the total glomeruli volume (GL) (6.7-fold from ∼3.2×10^6^μm^3^ to ∼21.5×10^6^μm^3^), and especially in the volume of the antennal lobe hub (ALH) (17-fold from ∼3.2×10^5^μm^3^ to ∼5.6×10^6^μm^3^). However, we did not find evidence for significant variation between *Heliconius* and non-*Heliconius,* in any of the four models tested (pMCMC>0.05) (Fig. 2), or between narrower phylogenetic groupings (i.e. outgroup genus or within *Heliconius* subclades) (pMCMC>0.05). Our analysis of phylogenetic signal suggests that the variation that we see for the three neuropils of interest (AL, λ=0.710; ALH, λ=0.671 and GL, λ=0.698), as well as the relationship between glomeruli and antennal lobe hub volumes (λ= 0.605), are only moderately influenced by phylogeny. Likelihood ratio testing comparing a model with and without phylogeny for all three neuropils and the fourth model showed that only the antennal lobe volume and the glomeruli volume have phylogenetic signal present (p<0.05). This suggest that the antennal lobe and glomeruli volumes are more strongly constrained by evolutionary history, whereas the antennal lobe hub volume and the relationship between glomeruli and antennal lobe hub volumes appear to be more influenced by other factors, such as ecological or functional constraints. In general, there were no interactions with sex detected in any of the four models (pMCMC>0.05), suggesting that the antennal lobe and antennal lob hub volumes of males and females are largely similar. However, we did find a steeper relationship between glomeruli volume and the antennal lobe hub volume in males than females (pMCMC=0.018).

Despite the lack of broad phylogenetic trends, pairwise comparisons revealed a number of differences between species (Table S1A-D). Due to sample size, these analyses were largely restricted to comparisons within *Heliconius* and some *Eueides* species for all 4 comparisons (i.e. AL, ALH, GL, and GL to ALH volume; Table S1A-D). Comparing the posterior distribution of each species’ random effects with each other, we noted that a few species constantly showed high probability that their posterior distribution is significantly different than the other species. This means that if the random effect for species A is greater, species A will have a higher baseline value for the trait bring measured compared to others. For example, *E. lybia* emerged as a clear outlier in the pairwise comparisons of individual antennal lobe structures, and in the scaling relationship between glomeruli and antennal lobe hub volumes (Fig. 2). We also noted that *E. lybia* had a greater total glomeruli volume than many other Heliconiini species. To test whether this increase in volume is associated with an increase in the number of glomeruli, we took a subset of five Heliconiini species, two *Heliconius,* two *Euiedes* and one *Dryadula* to examine if *E. lybia* had pronounced increases in glomeruli number. The number of glomeruli ranges from 65 to 87 (Table S2). Of the five species, *Dryadula phaetusa* had the greatest number of glomeruli (85-87), followed by *E. lybia* (82-83). However, *E. lybia* has the largest glomeruli volume (from ∼13×10^6^μm^3^ to ∼18.×10^6^μm^3^), and *Dryadula phaetusa* (from ∼10.8×10^6^μm^3^ to ∼12.6×10^6^μm^3^) had similar glomeruli volume to *H. erato* (from ∼11.5×10^6^μm^3^ to ∼12.4×10^6^μm^3^), despite the increased number of glomeruli (Table S2). *H. hecale* had the smallest number of glomeruli (68) and the smallest glomeruli volume (from ∼7.5×10^6^μm^3^ to ∼9.5.×10^6^μm^3^) (Table S2).

### Diversity of olfactory receptors across Heliconiini

As the number of glomeruli is thought to be directly related to the number of functional olfactory receptors, we fully annotated olfactory receptor genes (ORs) in 58 Heliconiini and 14 other Nymphalids genomes. In total we were able to annotate 4,658 genes (Table S3), which clustered into ∼70 orthologous groups (OGs) (Fig. 3), 21 of them specific to different subclades within Nymphalids. We recorded 10 distinct orthologous group duplications at the stem of Heliconinae (51, 40, 35a, 35b, 05, 04a, 04b, 04c, 56a and 56b), and five at the stem of Heliconiini (24, 24b, 05a, 05b and 56c) (Fig. 3, 4). Interestingly, OR05, belonging to the pheromone receptor clade, was duplicated once at the stem of Heliconinae and twice more at the stem of Heliconiini, while OR56, possibly linked to plant volatile compounds (Bastin-Héline et al. 2019), was found to be duplicated twice in Heliconinae and again, independently, in Heliconiini. As previously described (Cicconardi et al. 2023), we did not detect any duplications at the stem of *Heliconius*, also supporting previous findings that the great majority of duplications happened at base the of the tribe and subtribe radiation (Cicconardi et al. 2023). We also confirm three putative orthologous groups deletions of OR15, OR27, and OR35.

**Figure 3:**
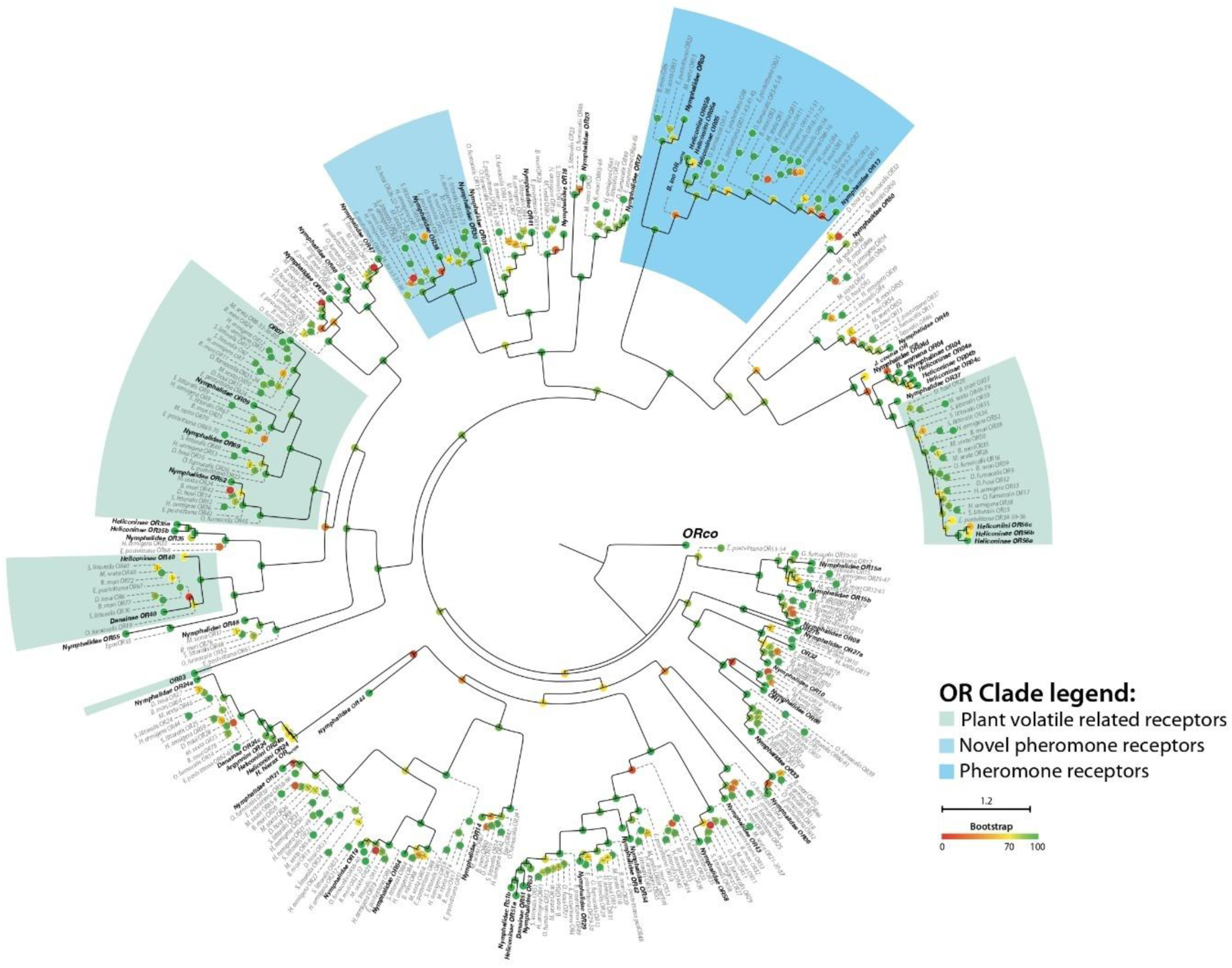
Approximate Maximum likelihood phylogeny reconstruction of olfactory receptor gene family across Lepidoptera using non-apilionoidea as reference (dashed lines). All monophyletic orthologous groups (OGs) are collapsed into a single node to show the entire phylogenetic tree, which contains 4658 butterfly genes and 428 reference genes. OGs containing butterfly sequences are listed in bold, together with its phylogenetic attribute. OGs without phylogenetic attribute are monophyletic at Lepidoptera level (e.g. ORco). The circle colours on nodes indicates bootstrap values.

**Figure 4:**
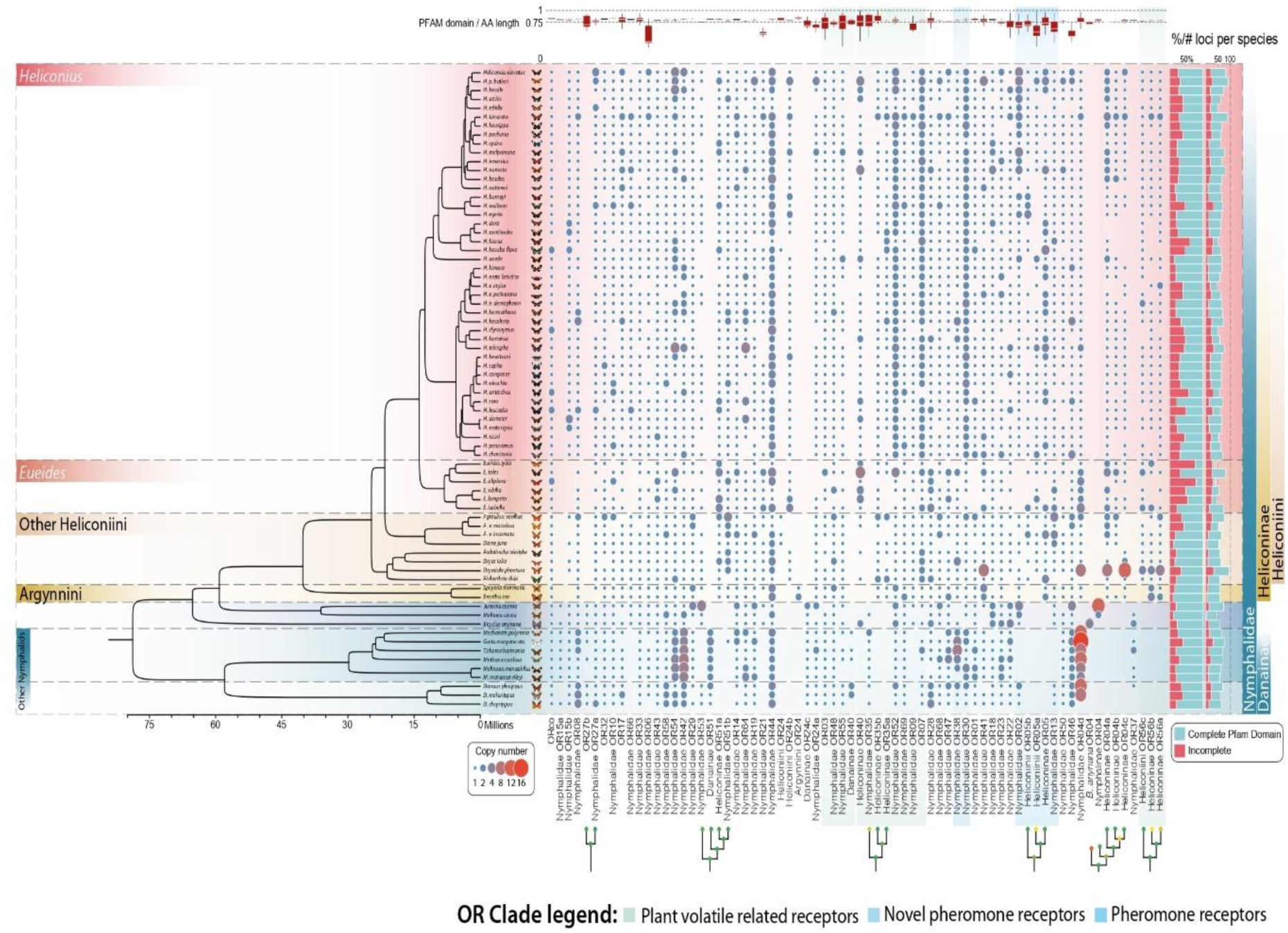
Copy number variation across Heliconinae and other Nymphalid species. On the left panel the ultrametric tree of all Nymphalid species generated with the chronos function (ape library), taking into account topologies and data phylogeny from (Cicconardi et al. 2023, 2024). In the central panel a bubble plot showing the expansions/contractions for each orthologous group (OG) (lower panel) for each nymphalid species. In the lower panel all butterfly OGs are listed with topological information for OG that are duplicated in butterflies. The right side of the panel shows a stacked histogram of percentage and absolute values of the number of OGs with complete or incomplete PFAM (7tm_6) domain. The upper panel shows instead boxplots of the distributions of fraction of protein sequence occupied by the 7tm_6 domain in each OGs. Broad distributions are a proxy of higher sequence variability.

Overall, *Heliconius* species have on average ∼65 ORs per species (excluding species with low quality assemblies), broadly within the range of other Heliconiini (and average 68 ORs) including *Agraulis* and *Dione* genera species (62 ORs). *H. aoedes*, a non-pollen feeding *Heliconius* species with 66 ORs, was also within range of other *Heliconius*. Despite this, within more closely related clades there are distinct outliers with higher numbers of ORs. For example, *H. pardalinus butleri* and *H. timareta* had 89 and 90 ORs, respectively, while *Eueides* species with the best quality genomes (*E. isabella* and *E. tales*) have 73 and 81 ORs, respectively. Of particular note is *Dryadula phaetusa*, which has 95 ORs, the largest repertoire within the whole dataset, which is in line with their high number of glomeruli. The OR expansion in *Dryadula* is driven largely by a large expansion of a few orthologous groups; OR04 (25 copies), OR41 (7 copies), and OR56 (from the pheromone clade; 11 copies).

Our MCMCglmm analyses, including the N50 value to account for genome completeness, did not show any significant relationships between OR number and the volumes of the three different neuropils (AL/ALH/GL) (Table S4). We obtained the same results when repeating the analyses with BUSCO completeness scores, instead of N50. Next, to explore whether the number of glomeruli is related to the number of olfactory receptors, we looked at the subset of data of four Heliconiini species in which all the individual glomeruli were segmented and counted. We observed that in three of the five species, the number of olfactory receptors was smaller than the number of glomeruli. However, two of these species (*E.lybia and E.aliphera*) had a BUSCO completeness score of less than 90%, which could have led to an underestimation of the number of olfactory receptors. Disregarding these 2 species, there was a disparity of ±5-10 between the olfactory receptor numbers and glomeruli numbers in the remaining 3 species (Table S2).

### Major environmental and ecological factors do not explain variation in antennal lobe morphology across the tribe

We found no significant association between any of the olfactory neuropils tested and the number of host plants used, the degree of social roosting, or the presence of pollen-feeding behaviours (Table S5). We also tested for associations with environmental factors that have been argued to affect the odorant concentration in the air, using datasets of annual temperature and annual precipitation. Once again, we did not find any significant associations between these variables and antennal lobe, antennal lobe hub, and total glomeruli volumes, or with the scaling relationship between the antennal lobe hub and total glomeruli volumes (Table S5). We further repeated these analyses with number of olfactory receptors as the independent variable, and again our results showed no significant associations (Table S5).

### Rearing environment influences relative volumes of the antennal lobe hub and glomeruli, but not overall antennal lobe volume

Our broad phylogenetic comparison of antennal lobe morphology took advantage of wild caught individuals. Although the differences observed might reflect heritable variation, they might also be the result of plastic responses to the environment. To test for potential effects of the local environment, we used a subset of five species from our wild dataset for which imaging was also available for insectary-reared individuals. We found that the interaction between species and origin (i.e. insectary-reared vs wild) was not significant in our comparison of antennal lobe volumes between wild and insectary individuals (χ2_4_=4.127, *P*_adj_=1.00), suggesting that the volumes between these two groups are similar across species (Table S6). There was no overall effect of origin on antennal volume (χ2_1_ =4.643, *P*_adj_=0.094), and only species (χ2_4_=12.341, *P*_adj_<0.05) and rCBR (as an allometric control) (χ2_1_ =14.628, *P*_adj_<0.05) were retained in our minimum adequate model (MAM). Subsequent analyses using SMATR on species-specific data showed no difference in the overall antennal lobe volume between wild and insectary individuals, with the exception of *H. melpomene* (significant slope shift; LR_1_=4.999, *P*_adj_<0.05), and *H. hecale* (significant grade shift; Wald χ2_1_ =4.505, *P*_adj_<0.05).

We next examined variation in the two constituent parts of the antennal lobe, the antennal lobe hub and total glomeruli volumes, and once again the interaction between species and origin was not significant (ALH: χ2_4_=3.453, *P*_adj_=1; GL: χ2_4_ =2.425, *P*_adj_=1.000). In both cases, species was not retained in the MAM, but there was a significant effect of origin (ALH: χ2_1_ =68.033, *P*_adj_<0.001; GL: χ2_1_ =41.297, *P*adj<0.001). Subsequent major axis regressions focused on the antennal lobe hub revealed that all differences that were observed among wild-caught and insectary-reared individuals in the five species tested were a result of significant grade shifts (α) or significant major-axis shifts, with the slopes being conserved (Table S7; Fig. S2). This pattern was similarly seen in the total glomeruli volume, with the exception of two species, *H. ismenius and H. hecale* (Table S7; Fig. S2).

Surprisingly, although we observed volumetric differences between the wild and insectary-reared individuals for both the antennal lobe hub and total glomeruli volume, these were not in the same direction. Specifically, although the antennal lobe hub was larger in insectary-reared individuals, the total glomeruli volume was smaller (Fig. 5). To investigate this relationship further we directly compared these two components of the antennal lobe. In our model of total glomeruli volume as a function of antennal lobe hub volume, neither the interaction between species and origin or species was significant (species*origin: χ2_4_=2.084, *P*_adj_=1.000, species: χ2_4_=11.435, *P*_adj_=0.066), suggesting that different species do not respond differently to captivity. However, there was an overall effect of origin (χ2_1_ =33.037, *P*_adj_<0.001). When looking at the scaling relationship between the total glomeruli volume and antennal lobe hub volume, we again observed a common slope, but an overall significant grade shifts across all five species (Fig. 5).

**Figure 5:**
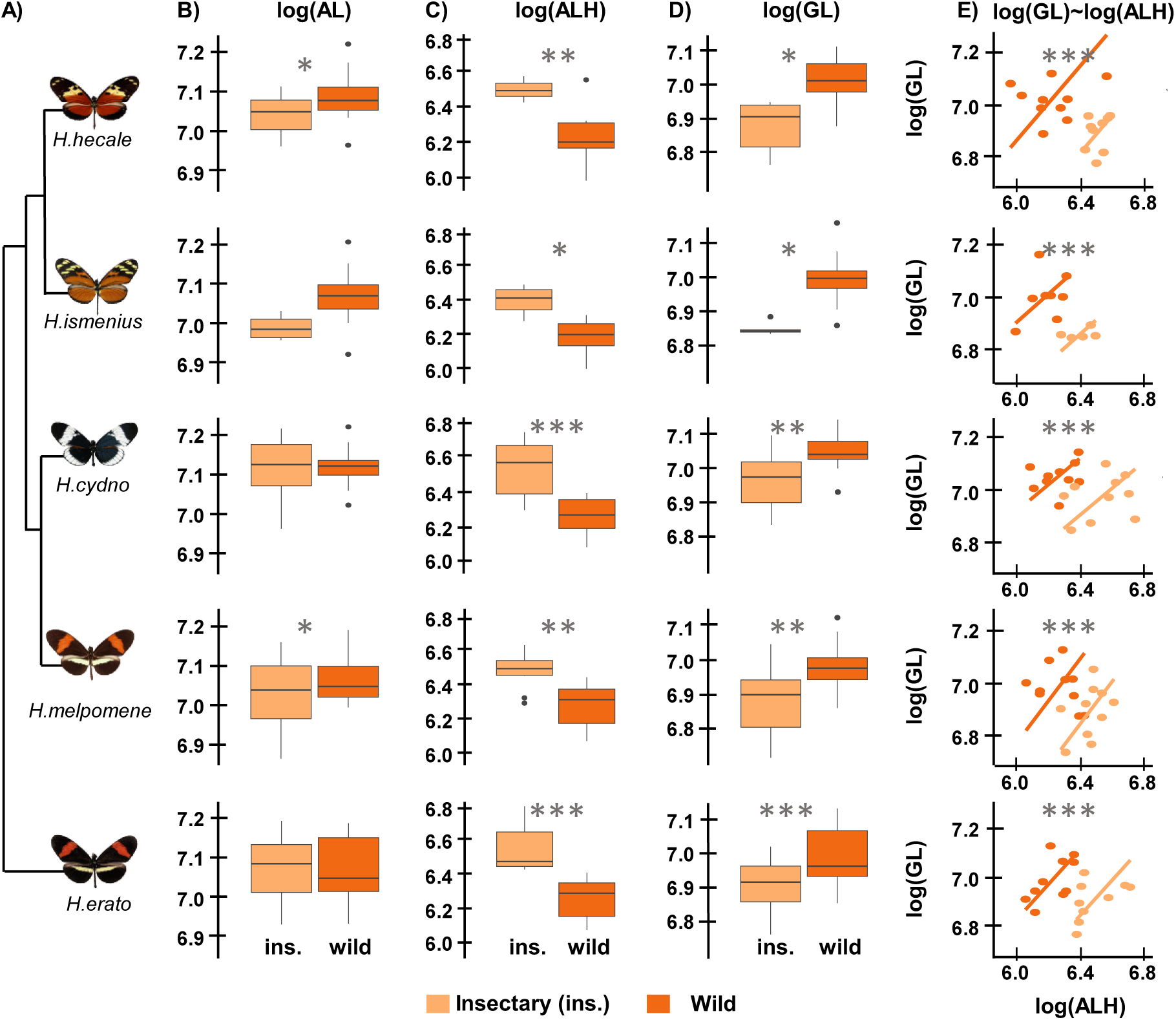
Comparisons of wild (dark orange) and insectary-reared (light orange) olfactory neuropil volumes for 5 species of *Heliconius*. **A)** phylogenetic relationship between *H. erato*, *H. melpomene*, *H. cydno*, *H. ismenius*, and *H. hecale*. **B), C)** and **D)** boxplots of the volume of AL, ALH, and GL respectively in insectary and wild individuals. **E)** Scaling relationship between GL and ALH volume in insectary and wild individuals across species. The common logarithm (log_10_) of all volumes were taken prior to analysis.

## Discussion

Groups of closely related, but ecologically diverse species provide an opportunity to test hypotheses about divergence in sensory and neural systems (Montgomery et al., 2021; Montgomery & Merrill, 2017; Wainwright & Montgomery, 2022). We examined variation in different components of the olfactory system across the Heliconiini tribe of Neotropical butterflies. Unlike previous studies in this group that have focused on a limited number of species pairs within this group (Montgomery et al., 2021; Montgomery & Merrill, 2017; Hebberecht et al., 2023), this allowed us to assess relationships between components of the olfactory system, and how these may correlate with ecological or environmental traits at a macroevolutionary scale. Our results reveal a mix of conserved and dynamic patterns of olfactory evolution, with species-specific shifts in antennal lobe morphology and olfactory receptor diversity occurring against a backdrop of broader phylogenetic stability. Notably, the variation in antennal lobe size across species appears to be driven by independent shifts in its two major components, the glomeruli and the antennal lobe hub, which likely reflect different selective pressures acting on these structures, and some independent variation between total glomeruli volume and OR number. We also uncovered a surprising pattern of developmental plasticity, with insectary-reared individuals exhibiting larger antennal lobe hubs but smaller glomeruli volumes compared to their wild-caught counterparts, suggesting complex environmental influences on the maturation of the olfactory system.

Although our results suggest that the architecture of the olfactory system is broadly conserved across the Heliconiini, we did observe substantial shifts in antennal lobe size for certain species. For example, *Eueides lybia* frequently emerged as an outlier in our analyses, displaying an overall increase in the antennal lobe, as well as significantly larger volumes of both its components, the glomeruli and antennal lobe hub volume. These findings might suggest enhanced olfactory discrimination capabilities linked to ecological demands, but current knowledge about *Eueides* provides little insight into the specific factors driving this difference. Similarly, for most species across the Heliconiini, we also see that investment into the glomeruli and the antennal lobe hub are mostly equal, again except for a few outlier species (e.g *Eueides lybia*), which suggest a general correlation of growth for the glomeruli and antennal lobe hub.

In addition to changes in the antennal lobe, we also found species-specific shifts in olfactory receptor number (e.g. *E. tales, E. isabella*, *H. timareta* etc.). *Dryadula phaetusa* exhibited the largest olfactory receptor repertoire in our dataset, driven by expansions in specific olfactory receptor subgroups, including those potentially linked to pheromone detection and plant volatile cues. Although further investigation of this interesting pattern is outside the scope of this study, we do also note that we distinguished ammino acid sequences having, lacking or partially lacking the PFAM domain for the 7-TMHs (7tm_6). Dryadula has 62 ORs with a complete conserved domain, in line with other Heliconiini, but has 28 loci with a shorter domain at the N-terminus. But only ∼10 of them are actually shorter than ∼300aa and have less than 5 TMHs; further analyses could evaluate the extent of which this lack of domain conservation is a limitation of the PFAM HMM or a signature of pseudogenisation.

Elsewhere, as with changes in antennal lobe volume, shifts in olfactory receptor numbers are difficult to ascribe to specific ecological events. The expansion of olfactory receptors in *H. timareta*, however, is intriguing given what we know of its ecology. Within a sympatric community, closely related *Heliconius* tend to differ in warning pattern, and it is well established that males often use colour to identify suitable mates (Merrill et al., 2011, 2015). This is certainly true for *H. cydno* (white or yellow) and its close relative *H. melpomene* (red), which coexist to the west of the Eastern Cordillera in the Andes (Merrill et al, 2019). However, on the eastern slopes of the Andes *H. cydno* is replaced by its sister species *H. timareta*, which shares the red patterns of the local *H. melpomene.* Interestingly, while neither species can discriminate based on colour alone, *H. timareta* (but not *H. melpomene*) males preferentially court *H. timareta* over *H. melpomene* females, presumably as a result of olfactory cues (Mérot et al., 2015; Rossi et al., 2024). As such, it is tempting to hypothesise that the expansion of olfactory receptors in *H. timareta* may have arisen as a result of a reinforcement-like process, to aid mate discrimination and avoid the production of unfit hybrids.

Overall, our analyses revealed little evidence to illuminate broader ecological correlates of species-specific expansions and reductions of the antennal lobe (and/or its constituent structures), or olfactory receptors. Broader trends in antennal lobe size evolution reported across the Lepidoptera have often revealed correlations with major shifts in activity pattern, and in particular diurnal versus nocturnal lifestyles (Stöckl et al., 2016; O’Donnell et al., 2013; Montgomery & Ott, 2015). In contrast, all Heliconiini species are diurnal, and *Heliconius* butterflies, in particular, are reported to predominantly forage for nectar and pollen in the early morning (Gilbert, 1975; Barp et al., 2011; Dell’Aglio et al., 2024b). Although we cannot discount undiscovered differences in diurnal activity across populations and species, evidence for this has not been found in captive populations (Dell’Aglio et al., 2024b). Although light intensity in closed forests will vary throughout the day, it will never be as low as at night, where light levels drop by eight orders of magnitude (Warrant, 2017). As such, it is perhaps unlikely that this difference necessitates significant investment in olfactory processing. Similar logic may also explain our failure to detect broad correlates between evolution in olfactory structures and ecology. Indeed, while shifts in antennal lobe volume have been reported among closely related *Heliconius* species inhabiting different forest types (Montgomery & Merrill, 2017), there is little evidence that this association is repeated across different species pairs, in contrast to the changes in the optic lobe (Montgomery et al., 2021, Rivas-Sánchez et al.; 2024). Indeed, across more diverse communities of ithomiine butterflies, variation in the visual system is more extensive and more closely tied to variation in habitat use than the olfactory system (Wainwright et al., 2024).

The species-specific variation we see in antennal lobe volume could stem from changes in the antennal lobe hub, or from changes in glomeruli volume. Overall, our analyses suggest that variation in antennal lobe size across species can result from both differential investment in glomeruli volume (6.7-fold absolute change) and antennal lobe hub volume (17-fold absolute change), however the range of variation in glomeruli volume is more similar to that for overall brain size (7.7-fold absolute change). While in some cases changes in the antennal lobe hub and glomeruli are correlated, this was not always the case, suggesting that these two components can evolve semi-independently. For example, *E. lybia* and *E. isabella* stood out with large glomeruli volumes, while other species like *H. erato cyrbia* and *H. himera* show disproportionately larger antennal lob hub volumes relative to glomeruli.

An increase in overall glomeruli volume can result from two factors, larger individual glomeruli or an increase in glomeruli number. Larger individual glomeruli would suggest an increase in olfactory receptor neurons housed within each glomerulus. Because studies have linked glomerular volume to olfactory receptor neuron number, this expansion may enhance sensitivity to olfactory cues (Takagi et al., 2024; Linz et al., 2013; Meisami, 1989), and *Drosophila* species with more but same types of olfactory receptor neurons have been shown to track host odors more efficiently and persistently (Takagi et al., 2024). Alternatively, glomeruli volume may also arise from an increase in glomeruli number, as suggested by our data on a subset of *Heliconiini* species (see below; Table S2). Since each glomerulus is expected to process input from a single class of olfactory receptor neuron (Hildebrand & Shepherd, 1997), an increased number of glomeruli could enhance odor discrimination by expanding the combinatorial possibilities for olfactory recognition. This has been observed in hornets, where the region of the antennal lobe with the most extensive variation (T_B_ Cluster) also had the highest glomeruli count, likely due to the greater demand for olfactory discrimination in high-density environments (Couto et al., 2021).

Changes in antennal lobe volume may also be driven by variation in antennal lobe hub volume. For instance, previous research found that differences in antennal lobe size between the glasswing butterfly (*Godyris zavaleta*) and two *Heliconius* species (*H. erato* and *H. hecale*) were largely due to expansion of the antennal lobe hub in *Godyris* rather than changes in glomeruli size (Montgomery et al., 2016). Since the antennal lobe hub contains axons of interneurons that integrate and relay olfactory information to integrative centres, its expansion could reflect differences in how olfactory signals are integrated across glomeruli and processed (Montgomery et al., 2016). In particular, a larger antennal lobe hub may enhance signal integration or prioritize specific odor types, potentially influencing behavioral responses.

The variation in glomeruli number in our dataset contrasts with the relatively constant glomeruli count observed across Lepidoptera, which typically ranges between 60 and 70 (Boeckh & Boeckh, 1979; Rospars, 1983; Berg et al., 2002; Kazawa et al., 2009; Carlsson et al., 2013), albeit with some exceptions, such as the oligophagous moth genus *Cydia*, which only has around 50 glomeruli (Varela et al., 2009; Couton et al., 2009; Trona et al., 2010). In contrast to this apparent consistency, we found that glomeruli number ranged from 65 to 87 in the five sampled Heliconiini species (Table S2). Although measurement errors and the lack of clear boundaries between individual glomeruli may have led to slight under- or overestimation, the number of glomeruli in our dataset aligns relatively well with the range of odor receptors identified in a much larger sample of Heliconiini. While the higher number of odor receptors compared to glomeruli in *Dryadula phaetusa* could be attributed to the expansion of certain odor receptor groups, we also acknowledge the possibility of functional redundancy or divergence in these butterflies. This could result in a deviation from the traditional 1:1 model of olfactory coding observed in other insects (Couto et al., 2005). However, future studies could expand the glomeruli counts to more species in the Heliconiini to get a more robust association between olfactory receptor number and glomeruli number, and single cell sequencing methods could provide insights into possible redundancy and co-expression of ORs in single sensory neurons.

Our analyses also reveal intriguing patterns of developmental plasticity. Insectary-reared individuals displayed larger antennal lobe hub volumes, but reduced glomeruli volumes compared to their wild-caught counterparts. This leads to a significant grade shift in the relationship between glomeruli and antennal lobe hub volumes between wild and insectary bred individuals. This divergent pattern suggests that the absence of natural environmental stimuli in captivity may differentially influence the development of these two structures. While the biological implications of these changes remain unclear, they raise questions about the potential trade-offs in sensory investment under constrained environments. It is possible that some of the interspecific variation we observe in our larger sampling of wild individuals reflect environmentally induced effects. However, the lack of current ecological correlates makes it unlikely that this would be driven by consistent environmental cues, although we cannot reject the possibility. However, examining olfactory receptor numbers in three species that differ greatly in terms of their antennal lobe hub (*H. himera*), glomeruli (*H. sara*) and glomeruli to antennal lobe hub volume (*H. aoede*) from other species — and accounting for genome completeness scores — we found that their olfactory receptor counts (*H. himera*: 64, *H. sara*: 70, *H. aoede*: 66) were within or close to the average for Heliconiini (68 ORs). This suggests that, despite their neuroanatomical differences, these species likely have similar capacities to process olfactory information, reducing the likelihood that their divergence is driven by evolutionary plasticity, although olfactory receptors turnover may result in different olfactory tuning. Expanding comparative studies of wild versus insectary-reared individuals to include species we identify as outliers would help eliminate the possibility of evolutionary plasticity. Additionally, experimental studies measuring olfactory sensitivity and behavioural outcomes in both wild and captive derived individuals could provide valuable insights into the functional consequences of the plastic responses in the glomeruli and antennal lobe hub.

Overall, our findings have important implications for future studies that use antennal lobe size as a proxy for olfactory capabilities across species. Our study demonstrates that glomeruli and antennal lobe hub volumes can evolve independently, with the environment acting on them in unexpected and even opposing ways. Consequently, total antennal lobe size might not be a straightforward indicator of olfactory abilities, underscoring the need to examine the functional contributions of glomeruli and antennal lobe hub separately in evolutionary and ecological contexts. In the Heliconiini, olfactory evolution is shaped by species-specific adaptations and developmental plasticity. Modular adjustments within the antennal lobe—for example, differential investment in glomeruli versus antennal lobe hubs— provide a flexible framework for rapid adaptation to diverse ecological niches. Although we did not observe direct associations with specific ecological traits, species-level differences, such as the large glomerular volume in *Eueides lybia* and the expansion of olfactory receptors in species like *H. timareta*, suggest that diverse evolutionary pressures are at play, warranting further investigation into unmeasured ecological or behavioural factors. Notably, our identification of a curious pattern of developmental plasticity—where insectary-reared individuals exhibited larger antennal lobe hubs but smaller glomeruli volumes compared to their wild-caught counterparts—highlights the influence of environmental conditions on olfactory system development and raises questions about how such plasticity may affect sensory processing and behaviour. Together, these results emphasize the importance of integrating evolutionary, ecological, and developmental perspectives when investigating olfactory evolution.

## Data availability

The underlying data and R-scripts supporting the findings of this study are available as part of the supplementary data “Heliconiini_olfactory_evolution_supplementary_tables.xlsx” and on github (https://github.com/SpeciationBehaviour/olfactory_system_evolution_heliconiini) respectively.

## Supporting information

FigureS1.tree

## Acknowledgements

We thank the environmental ministries of Costa Rica, Panama, French Guiana, Ecuador and Peru for permission to collect and export the sample in previous studies. We are grateful to the Wolfson Bioimaging Centre, University of Bristol, and the Center for Advanced Light Microscopy at the University of Munich Biocenter for imaging assistance. This work was supported by NERC Independent Research Fellowship NE/N014936/1 (SHM), and ERC Starter Grants 758508 (SHM) and 851040 (RMM).

## Role of authors

Conceptualisation: SHM, RMM; Data curation: YP, FC, SHM; Investigation and Methodology: YP, FC, SHM; Writing – original draft: YP, FC; Writing – review and editing; YP, FC, GB, RMM, SHM. Visualisation: YP, FC, GB; Funding acquisition, Project Administration and Supervision: SHM, RMM.

