## supplementary information for "Evolution of the olfactory system during the radiation of Heliconiini butterflies"

**This word document contains:**

**Supplementary Information 1:**

1.1 Table S2: Glomeruli counts, volume, receptor numbers and genome completeness scores of individuals from 4 species in the Heliconiini
1.2 Table S4: pMCMC values in our phylogenetically controlled comparative analyses with odor receptor numbers and its corresponding controls.
1.3 Table S5: pMCMC values in the phylogenetically controlled comparative analyses with additional environmental and ecological variables
1.4 Table S6: Statistical comparisons of different lmer models testing for species and origin significance
1.5 Figure S1: Approximate Maximum likelihood phylogeny reconstruction of the olfactory receptor gene family across Lepidoptera
1.6 Figure S2: Scaling relationships between wild and insectary-reared individuals of 5 *Heliconius* species

**Supplementary Information 2: Details of supplementary table included in 'Heliconiini_olfactory_evolution_supplementary_tables.xlsx':**

2.1 Table S1 A-D: MCMC pairwise comparisons of Heliconiini species
2.2 Table S3: Full annotation of 4658 olfactory receptor genes across Heliconiini
2.3 Table S7: SMATR statistics for comparisons of different olfactory neuropils between wild and insectary individuals in 5 *Heliconius* species
2.4 Table S8: Full volumetric, environmental and ecological data of wild Heliconiini samples
2.5 Table S9: Filtered dataset of Heliconiini samples for pairwise comparisons
2.6 Table S10: Full volumetric data of wild and insectary *Heliconius* species.

**Supplementary Information 1:**

**Table S2:** Table of glomeruli number, volume, odor receptor numbers, metrices for the assessment of genome completion (N50, BUSCO) and BUSCO completeness score for individuals from 4 species in the Heliconiini. OR stands for odor receptor. All glomeruli volume were $log_{10}$ before analysis.

| Species | Sex | Glomeruli number | Glomeruli volume ($\mu m^3$) | OR numbers | N50 | BUSCO | BUSCO score |
|---|---|---|---|---|---|---|---|
| *E. lybia* | Male | 83 | $18.x10^6$ | 35 | 4997 | 951 | 58.1 |
| *E. lybia* | Male | 82 | $13x10^6$ | | | | |
| *E. aliphera* | Female | 65 | $7.6x10^6$ | 38 | 2835 | 569 | 34.5 |
| *E. aliphera* | Male | 77 | $8.5x10^6$ | | | | |
| *E. aliphera* | Female | 75 | $7.9x10^6$ | | | | |
| *H. erato demophoon* | Male | 69 | $11x10^6$ | 65 | 10689 | 1617 | 98.7 |
| *H. erato demophoon* | Male | 70 | $12x10^6$ | | | | |
| *H. hecale* | Male | 68 | $7.5x10^6$ | 73 | 93 | 1609 | 97.8 |
| *H. hecale* | Female | 68 | $9.5.x10^6$ | | | | |
| *D. phaetusa* | Female | 87 | $10.8x10^6$ | 95 | 6136 | 1658 | 100 |
| *D. phaetusa* | Female | 85 | $12.6x10^6$ | | | | |

**Table S4:** Table of pMCMC values in our phylogenetically controlled comparative analyses with odour receptor numbers and its corresponding controls. OR variables in all analyses had non-significant associations (pMCMC>0.05). Antennal lobe= AL; Antennal lobe hub= ALH; Glomeruli= GL; OR= odor receptor. All numbers rounded to 3d.p.

| | Dependent Variables | | | |
|---|---|---|---|---|
| **Neuropils** | **OR+BUSCO** pMCMC (OR) | **OR+BUSCO** pMCMC (BUSCO) | **OR+N50** pMCMC (OR) | **OR+N50** pMCMC (N50) |
| AL | 0.461 | 0.145 | 0.947 | *0.041 |
| ALH | 0.335 | 0.614 | 0.337 | 0.184 |
| GL | 0.506 | 0.088 | 0.882 | 0.131 |

**Table S5:** Table of pMCMC values in our phylogenetically controlled comparative analyses with additional environmental and ecological variables. All variables had non-significant associations (pMCMC>0.05). Antennal lobe= AL; Antennal lobe hub= ALH; Glomeruli= GL; OR= odor receptor. All numbers rounded to 3d.p.

| | Neuropils | | | | | |
|---|---|---|---|---|---|---|
| | **AL~rCBR** | **ALH~rCBR** | **GL~rCBR** | **GL~ALH** | **OR~N50** | **OR~BUSCO** |
| **Variables** | pMCMC | pMCMC | pMCMC | pMCMC | pMCMC | pMCMC |
| annual temperature | 0.380 | 0.567 | 0.224 | 0.865 | 0.643 | 0.594 |
| annual precipitation | 0.563 | 0.251 | 0.104 | 0.176 | 0.908 | 0.298 |
| relative humidity | 0.573 | 0.924 | 0.431 | 0.492 | 0.539 | 0.447 |
| windspeed | 0.300 | 0.198 | 0.824 | 0.173 | 0.371 | 0.763 |
| presence of pollen-feeding | 0.324 | 0.429 | 0.586 | 0.492 | 0.765 | 0.988 |
| number of host plant use | 0.553 | 0.627 | 0.798 | 0.306 | 0.633 | 0.496 |
| degree of social roosting | degree 1: 0.551<br>degree 2: 0.424<br>degree 3: 0.559 | degree 1: 0.298<br>degree 2: 0.565<br>degree 3: 0.541 | degree 1: 0.863<br>degree 2: 0.610<br>degree 3: 0.604 | degree 1: 0.482<br>degree 2: 0.782<br>degree 3: 0.537 | degree 1: 0.267<br>degree 2: 0.988<br>degree 3: 0.710 | degree 1: 0.855<br>degree 2: 0.390<br>degree 3: 0.961 |

**Table S6:** Comparisons of different lmer models. **A**) Values from Antennal lobe (AL), Antennal lobe hub (ALH) and Glomeruli (GL) volumes, and **B**) GL~ALH volume comparisons across all individuals, testing for significance of including species and origin in the model. Reduced models indicate that the variables were excluded from the full model stated. Volume of the rest of the Central Brain (rCBR) was used as the allometric control. All neuropil volume were $log_{10}$ before analysis.

**A**) comparisons across all individuals: testing for species and origin effect
wild+insectary individuals: 5 species (n=92)

| Neuropil | Reduced models | χ2 | Df | p | adjusted p | Full model |
|---|---|---|---|---|---|---|
| AL | -* | 4.13 | 4 | 0.39 | 1 | lmer (Neuropil~rCBR+sp*wild/insectary + (1\|sex)) |
| | -* -sp | 12.34 | 4 | 0.01* | 0.04* | |
| | -* -wild/ insectary | 4.64 | 1 | 0.03* | 0.09 | |
| ALH | -* | 3.45 | 4 | 0.49 | 1 | |
| | -* -sp | 7.19 | 4 | 0.13 | 0.38 | |
| | -* -wild/ insectary | 68.03 | 1 | < 0.001 *** | <0.001 *** | |
| GL | -* | 2.42 | 4 | 0.66 | 1 | |
| | -* -sp | 10.11 | 4 | 0.04 * | 0.07 | |
| | -* -wild/ insectary | 41.30 | 1 | <0.001 *** | <0.001 *** | |

**B**) comparisons across all individuals for rate of change between GL and ALH:
testing for species and origin effect

| Reduced models | χ2 | Df | p | adjusted p | Full model |
|---|---|---|---|---|---|
| -* | 2.08 | 4 | 0.72 | 1 | lmer(GL~ALH+sp*wild/insectary +(1\|sex)) |
| -* -sp | 11.44 | 4 | 0.02* | 0.07 | |
| -* -wild/ insectary | 33.04 | 1 | <0.001 *** | <0.001 *** | |

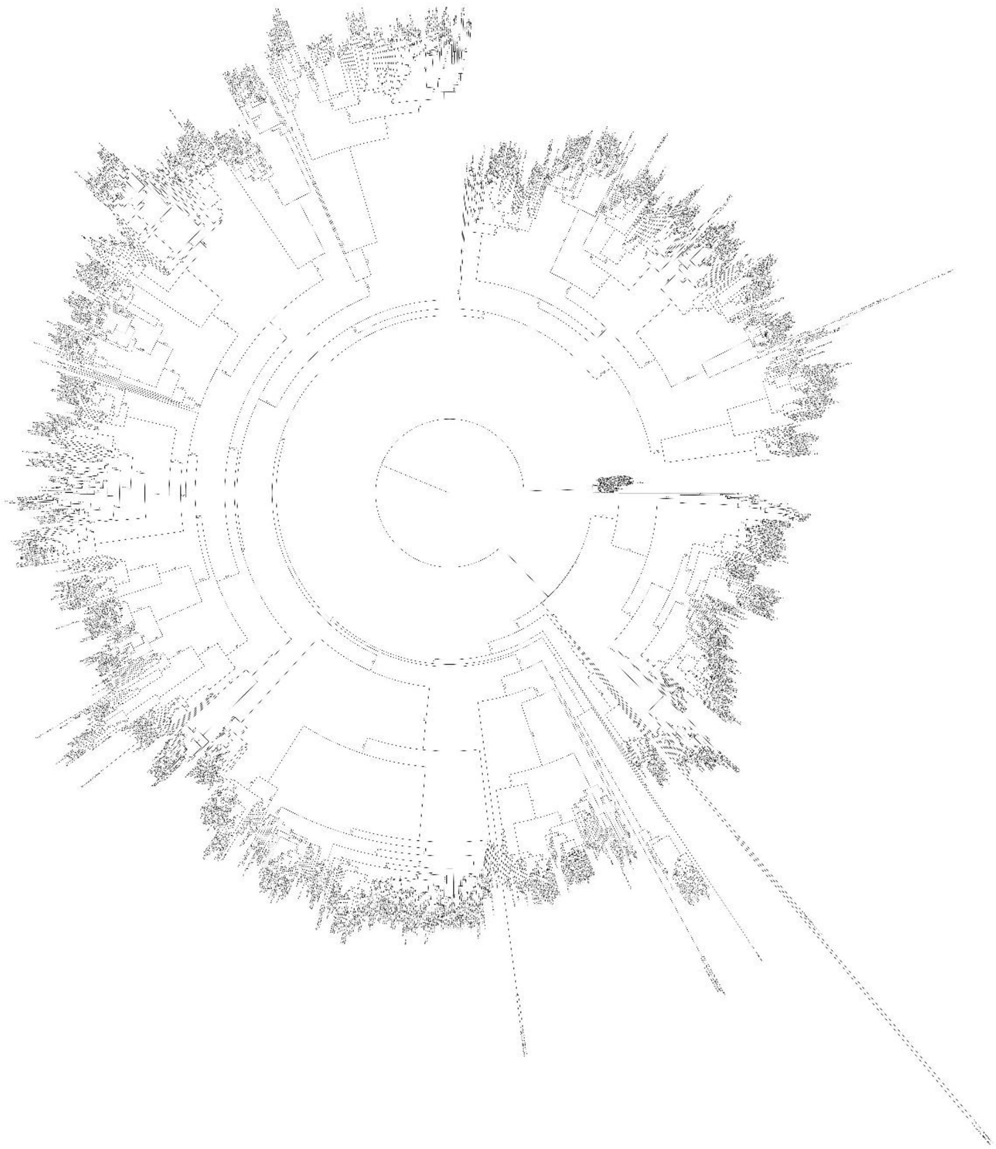

**Figure S1: Approximate Maximum likelihood phylogeny reconstruction of the olfactory receptor gene family across Lepidoptera using non-Papilionoidea as reference, rooted using the ORco OG. A TREES.file of this phylogeny reconstruction is also available as additional supplementary materials.**

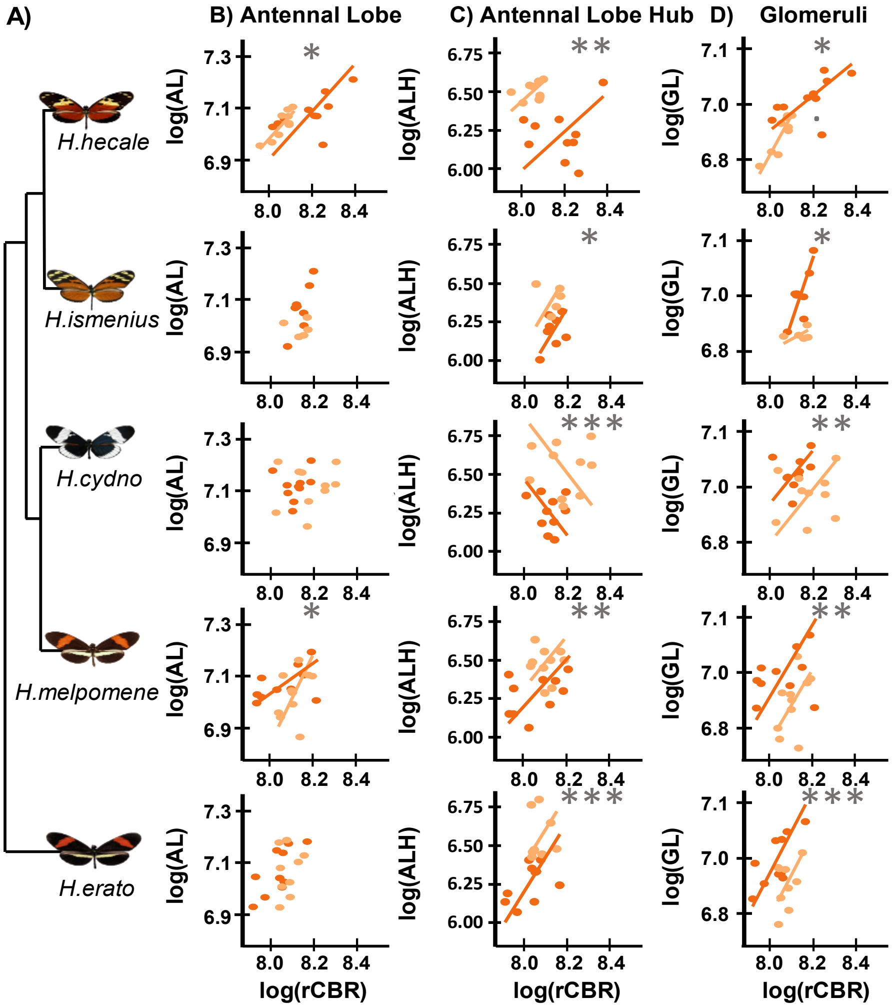


**Figure S2: Within species comparisons of the scaling relationships amongst different neuropils of interest between wild (dark orange) and insectary-reared (light orange) individuals across 5 species of Heliconius**. **A)** phylogenetic relationship between *H.erato, H.melpomene, H.cydno, H.ismenius*, and *H.hecale*. **B, C,** and **D)** Scaling relationship between the rest of Central Brain (rCBR) and total Antennal lobe (AL), Antennal lobe hub (ALH), and Glomeruli (GL) volume respectively in insectary and wild individuals across species. The common logarithm (log10) of all volumes were taken prior to analysis.

### Supplementary Information 2: Details of data included in 'Heliconiini_olfactory_evolution_supplementary_tables.xlsx'

**Table S1: MCMC pairwise comparisons of the volumes of A)** the antennal lobe, **B)** antennal lobe hub, **C)** glomeruli and **D)** glomeruli~antennal lobe hub in the Heliconiini. Only species with more than 5 individuals are included in the final comparison table. Final row in all tables indicates the number of pairwise differences cumulated for each species, for each region. Numbers in other cells indicates the difference in posterior distribution of the random effect of one species compared to the other species. Numbers close to 0 indicates that the species in the row consistently has a smaller random effect than the species in the column while numbers close to 1 indicates that the species in the row consistently has a larger random effect than the species in the column. *** represent a significant posterior probability (more than 99% or less than 1%) that 2 species differ in their neuropil volumes and are highlighted in bold. Top 3 species with the greatest number of differences are similarly highlighted in bold. Species names are abbreviated. All comparative numbers are rounded up to 3d.p. All neuropil volume were $\log_{10}$ before analysis.

**Table S3: Full annotation of 4658 olfactory receptor genes across Heliconiini.** The table describes the annotated OR loci across all Nymphalid species taken into account. It also describes some of the features of the sequences and loci. The column names from bed1 to bed12 correspond to the full genomic coordinates of these genes.

**Table S7: SMATR statistics for comparisons of different olfactory neuropils between wild and insectary individuals in 5 *Heliconius* species**. A significant adjusted p-value indicates a significant difference in that particular neuropil between wild and insectary individuals. All neuropil volume were $\log_{10}$ before analysis. Antennal lobe= AL; Antennal lobe hub= ALH; Glomeruli= GL and rest of Central Brain=rCBR. All numbers rounded to 3d.p.

**Table S8: Full volumetric, environmental and ecological data of wild Heliconiini samples.** sim.clade= simplified clade groupings, sp abb= species abbreviation. AL=antennal lobe, ALH=antennal lobe hub, Gl= glomeruli, rCBR= rest of central brain. All neuropil volumes are in $\mu m^3$ . AT= annual temperature, AP= annual precipitation, RH= relative humidity, PF= presence of pollen feeding, with 0 indicating no and 1 indicating yes. SR= social of degree roosting, No.HP= number of host plants used, OR= number of odour receptors.

**Table S9: Filtered dataset of Heliconiini samples for pairwise comparisons.** A subset of samples from Table S8. Only species that have more than or equal to 5 individuals are included. sim.clade= simplified clade groupings, sp abb= species abbreviation. AL=antennal lobe, ALH=antennal lobe hub, Gl= glomeruli, rCBR= rest of central brain. All neuropil volumes are in $\mu m^3$ .

**Table S10: Full volumetric data of wild and insectary *Heliconius* species**. A dataset of samples from 5 Heliconius species with data of individuals from the wild and the insectary. sim.clade= simplified clade groupings, sp abb= species abbreviation. AL=antennal lobe, ALH=antennal lobe hub, Gl= glomeruli, rCBR= rest of central brain. All neuropil volumes are in $\mu m^3$ .
