## Supplementary figures and images for "Evolution of the olfactory system during the radiation of Heliconiini butterflies"

### FigureS1.tree

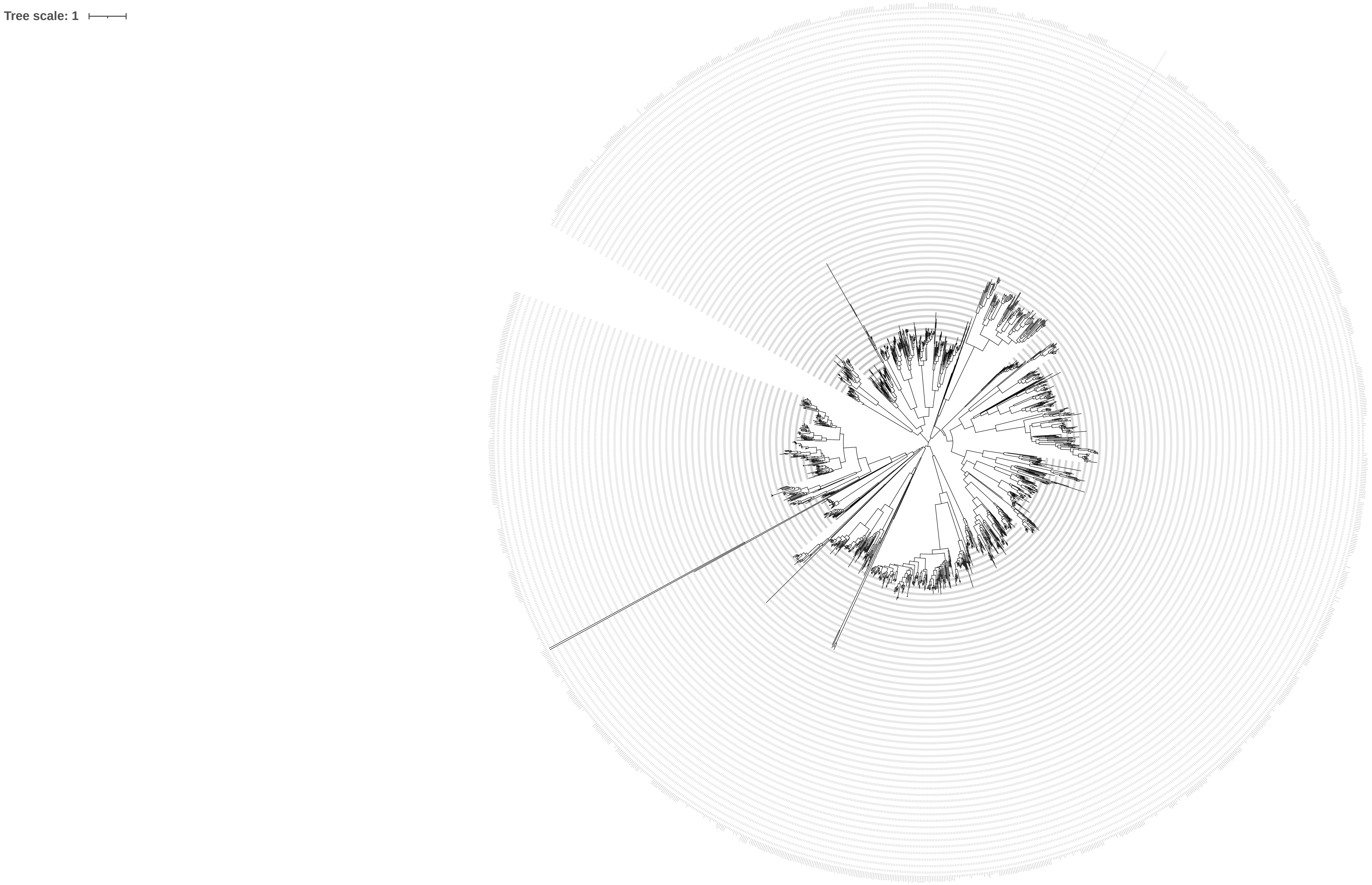
